# The cellular coding of temperature in the mammalian cortex

**DOI:** 10.1101/2022.02.21.481299

**Authors:** M. Vestergaard, M. Carta, J.F.A. Poulet

## Abstract

Temperature is a fundamental sensory modality separate from touch, with dedicated receptor channels and primary afferent neurons for cool and warm (Blix, 1882; Filingeri, 2016; Vriens et al., 2014). Unlike other modalities, however, the cortical encoding of temperature remains mysterious, with very few cortical neurons reported that respond to non-painful temperature and the presence of a ‘thermal cortex’ is debated (Bokiniec et al., 2018; Craig et al., 2000; Hellon et al., 1973; Milenkovic et al., 2014; Tsuboi et al., 1993). Using widefield and two-photon calcium imaging in the mouse forepaw system, here we identify cortical neurons that respond to cooling and/or warming with distinct spatial and temporal response properties. Surprisingly, we observed a representation of cool, but not warm, in the primary somatosensory cortex, but cool and warm in the posterior insular cortex (pIC). The representation of thermal information in pIC is robust, somatotopically-arranged and reversible manipulations show a profound impact on thermal perception. Intriguingly, despite being positioned along the same one-dimensional sensory axis, the encoding of cool and warm is distinct, both in highly- and broadly- tuned neurons. Together, our results show that pIC contains the primary cortical representation of skin temperature and may help explain how the thermal system generates sensations of cool and warm.

---

A fundamental question in neuroscience is: how is the external sensory environment represented in the cortex? In the thermal system, there is currently no consensus on how or where sensory information is encoded in the cortex. One model is that cool and warm are processed by functionally and anatomically segregated circuits, following labelled line principles seen in primary sensory afferent neurons and spinal circuits (Andrew and Craig, 2001; Craig et al., 2001; Darian-Smith et al., 1973, 1979; Duclaux and Kenshalo DR, 1980; Kenshalo and Duclaux, 1977; Ran et al., 2016; Wang et al., 2018; Yarmolinsky et al., 2016) resulting in cool and warm selective cortical neurons (Fig. 1a - top). Another is that cool and warm are integrated in the thermal pathway resulting in cortical neurons with a continuous, graded representation of temperature (Fig. 1a - bottom). Testing these models requires the identification of thermally responsive neurons in the cortex.

**Figure 1.**
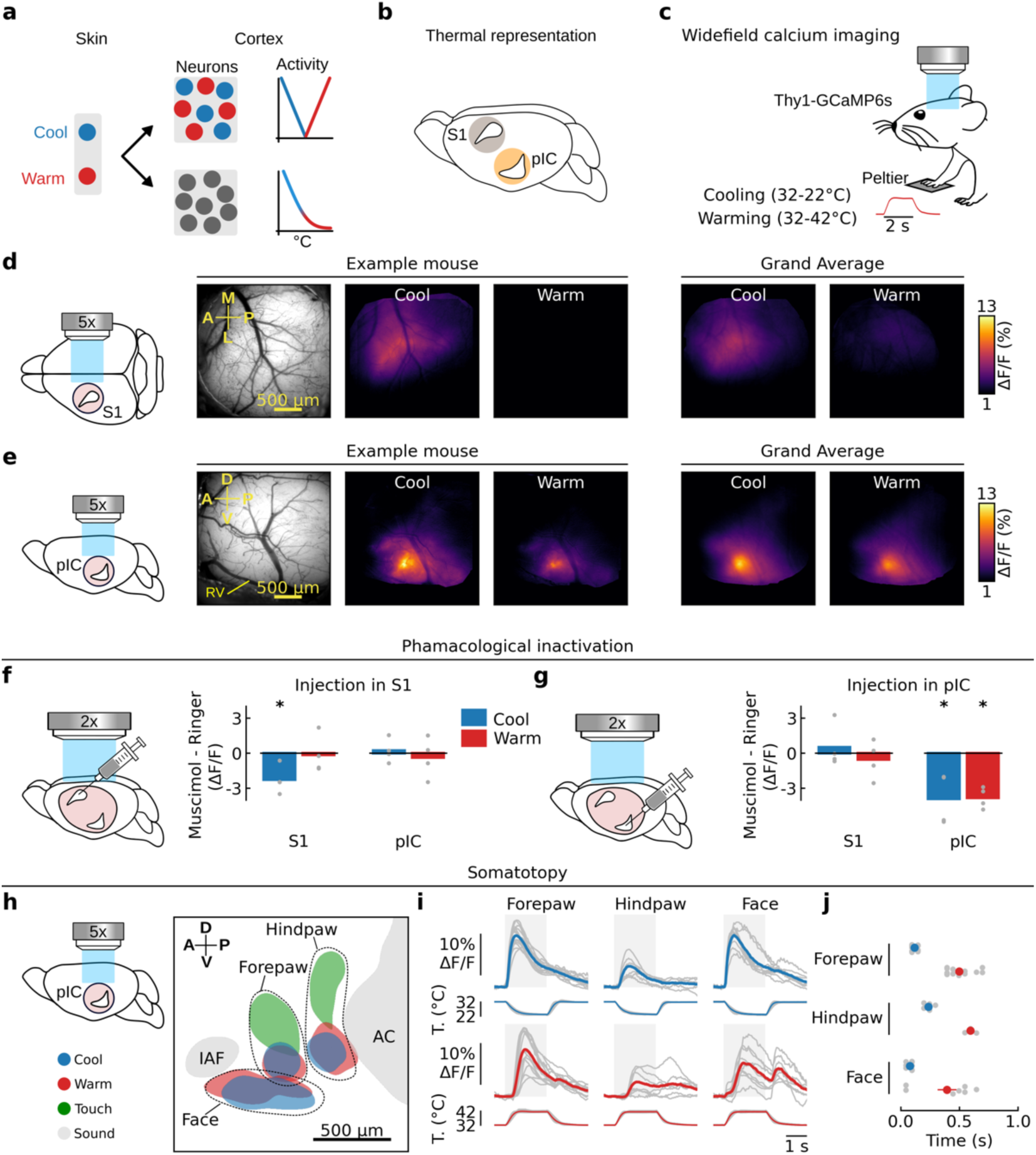
Cortical representation of cool and warm. **a**, Cartoon schematic showing segregated (top) or integrated (bottom) model of cortical thermal encoding. **b**, Mouse brain showing candidate locations of thermal cortex in primary somatosensory cortex (S1) or posterior insular cortex (pIC). **c**, Schematic of an awake Thy1-GCaMP6s mouse with right forepaw on a Peltier element during widefield calcium imaging, inset shows temporal dynamics of warming stimulus. **d**, From left to right, schematic of glass window on S1 (pink circle); in vivo image of cortical surface; averaged widefield response to 10°C cooling (32-22°C) or warming (32-42°C) of the forepaw in an example mouse; grand average across mice (*n* = 286 cool and 285 warm trials, 7 mice). M = medial, A = anterior, L = lateral, P = posterior. **e**, Same as **d**, but for mice with glass window implanted over pIC (*n* = 360 cool and 352 warm trials, 14 mice), RV = rhinal vein. D = dorsal, A = anterior, V = ventral, P = posterior. **f**, From left to right, schematic illustrating injection site into S1 during widefield imaging through a clear skull preparation with simultaneous imaging of S1 and pIC (pink shows field of view); bar graphs show difference in response amplitude following muscimol vs. Ringer’s injection. Bars indicate mean and gray filled circles indicate individual mice (*n* = 4). * shows significant difference in S1 response following muscimol vs Ringer’s injection (*p* < 0.05, two-sided *t*-test). **g**, Same as **f** but showing a reduction in pIC response to thermal stimuli after pIC inactivation and no change in the S1 response (*n* = 4 mice). **h**, Somatotopic map of responses locations to thermal (10°C cool and warm), tactile (100 Hz) and acoustic (8 kHz) stimulation. Colored area indicates peak population response averaged across mice (see methods, *n* = 14 thermal forepaw, 7 thermal hindpaw, 9 thermal face, 9 touch forepaw, 6 touch hindpaw, 13 sound). Data from individual mice are aligned to peak activity of the thermal forepaw response. IAF, insular auditory field; AC, auditory cortex. **i**, Widefield responses to thermal stimulation of different body parts from the same dataset as in **h**. Gray lines show mean responses from individual mice (*n* values same as **h**), colored lines show population mean, and grey area indicates time from start of stimulus to end of plateau phase. **j**, Gray filled circles show response latencies from individual mice, colored filled circles show mean ± s.e.m.

Psychophysical studies have shown that cooling and warming elicit distinct sensations, influenced by stimulus amplitude, duration, dynamics, body part and adapted skin temperature (Filingeri, 2016; Paricio-Montesinos et al., 2020). Analogous to other modalities, these features should be represented in a cortical region dedicated to thermal processing. Moreover, manipulation of its activity should influence thermal perception. A number of cortical regions have been suggested to be involved in thermal processing, including the primary somatosensory cortex (S1) (Hellon et al., 1973; Milenkovic et al., 2014; Tsuboi et al., 1993) and the posterior insular cortex (pIC) (Beukema et al., 2018; Craig et al., 2000; Penfield and Faulk, 1955) (Fig. 1b). However, no study has identified a cortical area with a cellular representation of cool and warm and a reversible impact on perception.

## Posterior insular cortex encodes somatotopically cool and warm, S1 only cool

To examine thermal processing in S1 and pIC, we performed widefield calcium imaging through a glass window in paw-tethered, awake mice expressing calcium indicator GCaMP6s in cortical excitatory neurons (Fig. 1b, c, Supplementary Fig. 1). We presented an 8 kHz acoustic stimulus to locate the anterior auditory fields (AAF) as well as the small diameter auditory insular field (IAF) that boarder the somatosensory regions of pIC (Gogolla et al., 2014; Rodgers et al., 2008; Sawatari et al., 2011), and went on to perform posthoc histology (Supplementary Fig. 1). Next, we delivered 2 s thermal stimuli to the forepaw glabrous skin via a Peltier element held at an adapted temperature (AT) of 32°C. Surprisingly, while 10°C (32°C to 22°C) cooling stimuli triggered reliable changes in fluorescence in S1, 10°C warming (32°C to 42°C) did not (Fig. 1d). In contrast, both cooling and warming stimuli triggered robust, large amplitude responses in pIC (Fig. 1e). Local pharmacological inactivation of either S1 or pIC during imaging abolished the thermal response in the injected region but not in the untreated region (Fig. 1f, g), indicating that pIC and S1 receive parallel streams of thermal input.

Perceptual thresholds for cool and warm covary across body parts (C. Stevens Kenneth K. Choo, 1998), suggesting that some body parts have a stronger cortical representation than others. Widefield calcium imaging showed a clear somatotopic arrangement of thermal and tactile responses anterior to the AAF in all mice (Fig. 1h) with the face represented in a region anterior and ventral to the forepaw which, in turn, is anterior to the hindpaw (Fig. 1h). In all mice, the forepaw had a larger amplitude response to thermal stimuli than the hindpaw, perhaps reflecting the dominant role of the forepaw in haptic exploration (Fig. 1h and i). Intriguingly, while responses to cooling and touch spatially overlap in S1 (Supplementary Fig. 2) (Milenkovic et al., 2014), they were separate in pIC (Fig. 1h). For all body parts tested, warming responses were delayed compared to cooling, suggesting that neural response latency is a hardwired property of the thermal system (Fig. 1j).

### Heterogeneous arrangement of thermally-tuned neurons in the pIC

Cooling and warming widefield responses in pIC overlap spatially (Fig. 1e, h). This could result from an intermingled distribution of highly tuned or broadly tuned cells (Fig 2a). To determine the thermal tuning of cortical neurons, we went on to perform two-photon calcium imaging of pIC excitatory neurons in awake mice and observed robust cellular responses to cooling and warming (Fig 2b, c, d). We went on to plot the normalized responses according to a thermal bias index that describes the relative response strength to cooling compared to warming with a value of -1 indicating a cool only cell and +1 warm only (Fig. 2d, e). The index had a U-shaped distribution, suggesting segregated channels of tuned input drive cool and warm responses, and showed a similar probability of highly tuned neurons (cool only or warm only) and broadly tuned (cool and warm responsive) neurons. In line with widefield data, the vast majority of neurons in S1 were tuned to cooling only and only a tiny fraction responded to warming with delayed and inconsistent responses (Fig. 2e, 3c and Supplementary Fig. 3).

**Figure 2.**
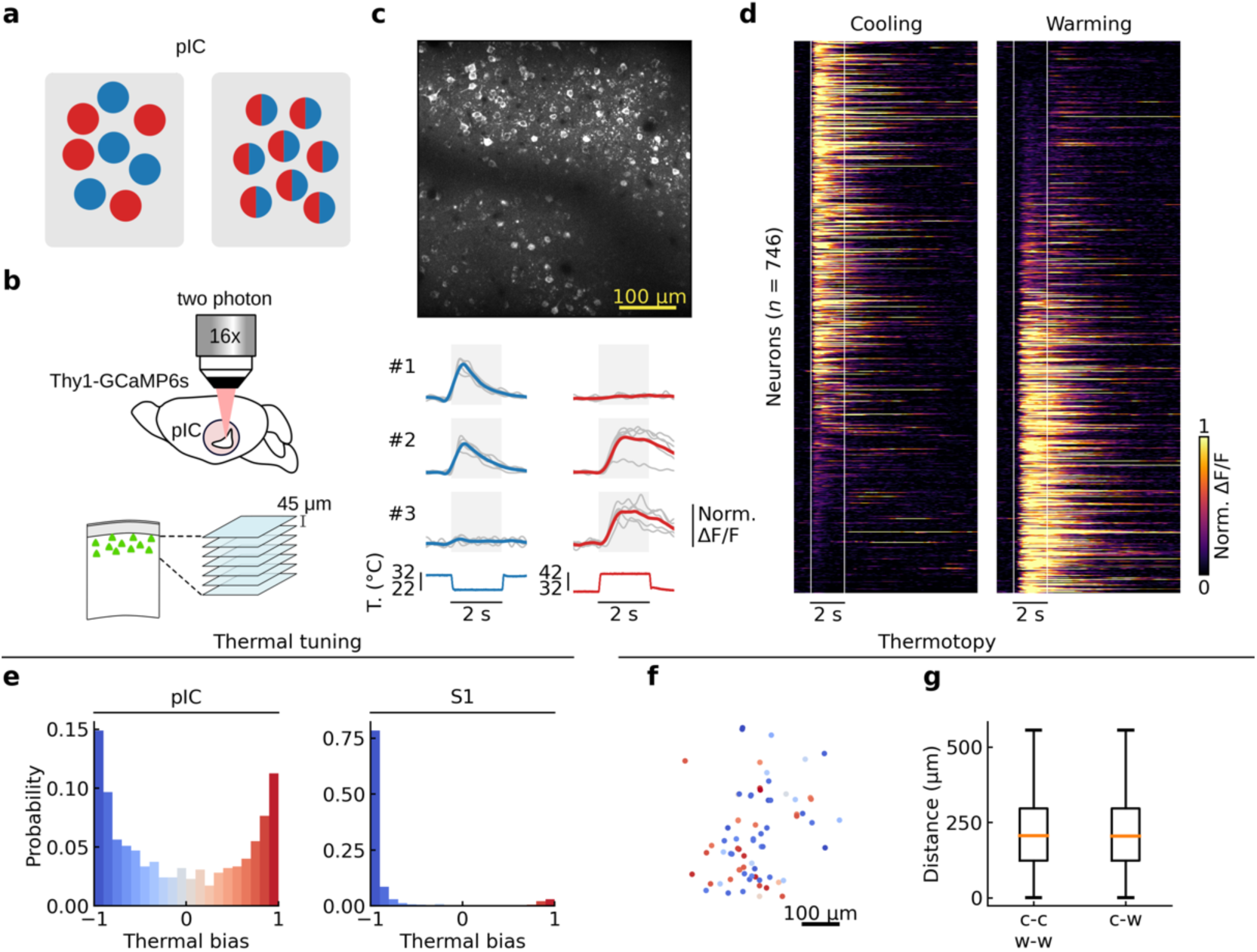
Heterogeneous arrangement of thermally-tuned neurons in the posterior insular cortex. **a**, Schematic showing (left) tuned vs. (right) broadly-tuned cortical cells. **b**, Schematic showing two-photon calcium imaging of pIC. Imaging started at 100 μm from the pial surface of a Thy1-GCaMP6s mouse, 7 optical sections were acquired with intervals of 45 μm. **c**, Top, example in vivo two photon image from pIC. Bottom, example of responses of single pIC neurons during 10°C cooling (blue) or 10°C warming (red) stimuli from an AT of 32°C (*n* = 5 trials each). Here and for all two photon data figures, gray lines show single trial responses, colored lines show average, grey area indicates time from start of stimulus to end of plateau phase. Below, corresponding stimulus traces. **d**, Single cell pIC calcium responses to cooling (left) and warming (right) stimuli. Each line represents a single neuron, responses are normalized to the peak and sorted based on the thermal bias index with vertical white lines showing the onset and the end of the plateau phase of thermal stimuli (*n* = 746 cells, 7 mice, 16 sessions). **e**, Histograms showing the distribution of thermal bias for all responsive cells in pIC (left) and S1 (right)(*n* = 411 cells, 4 mice, 9 sessions) **f**, Spatial map of neurons color coded with their thermal bias for the representative mouse shown in **c**. **g**, No significant difference in the distance between pairs of cells with similar bias (cold to cold (c-c) or warm to warm (w-w)) and distance between pairs of cells with opposite bias (cold to warm (c-w)) indicates no thermotopy for strongly biased cells (difference of medians confidence interval (−2 µm, 5 µm), 95%, bootstrapped).

**Figure 3.**
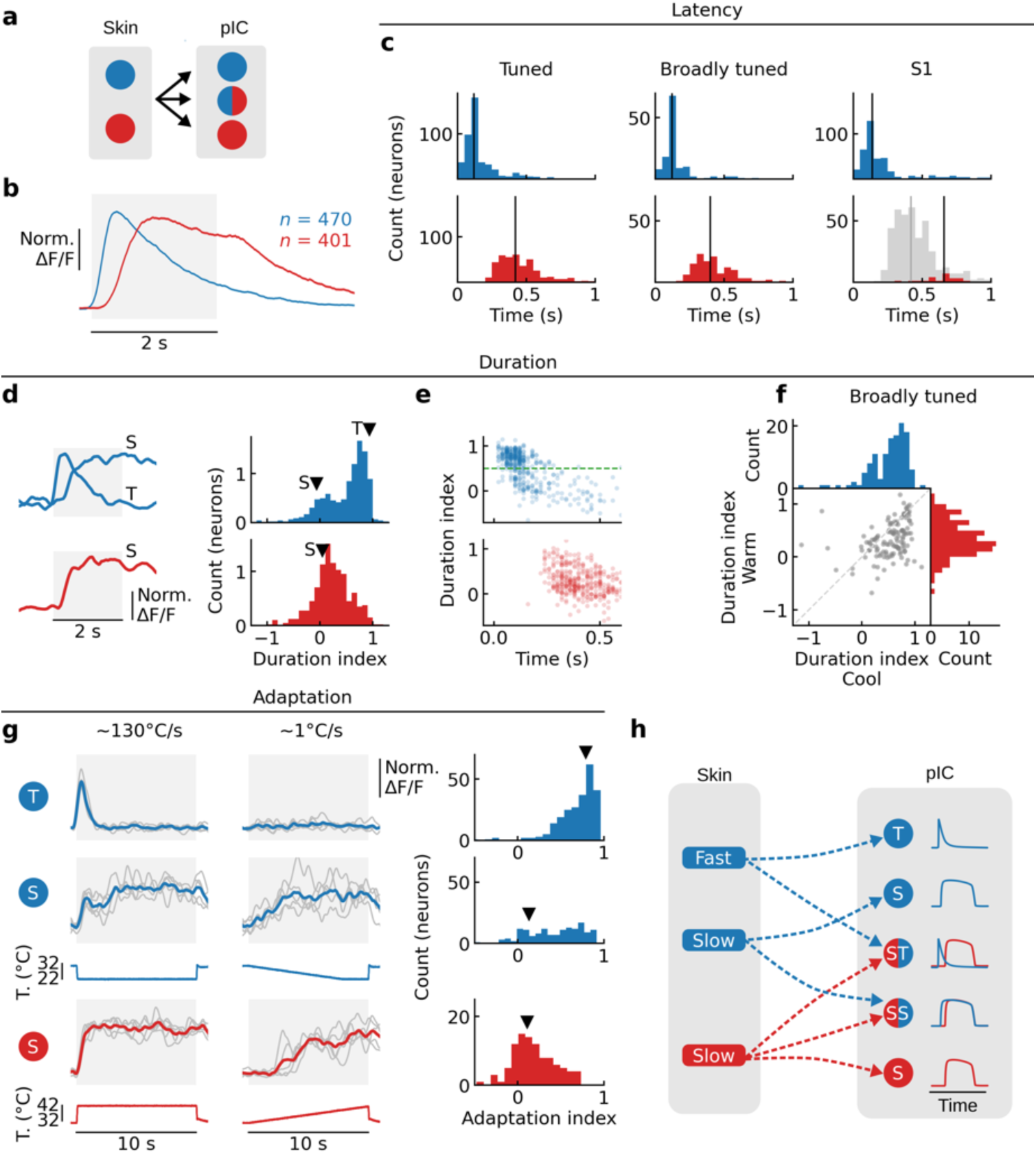
Distinct temporal dynamics of cool and warm encoding. **a**, Schematic showing transition from selective cool and warm skin spots to tuned and broadly tuned pIC neurons. **b**, Grand average responses (mean ± s.e.m.) to 10°C cooling and warming responses from AT of 32°C (*n* = 470 cool cells, 401 warm cells; these cells are also used in panels c, d and e. Dataset is the same presented in Fig 2). **c**, Histograms of the response latency for (top) 10°C cooling, and (bottom) 10°C warming, in (left to right) tuned, broadly tuned, and in S1 (grey bars show comparison to tuned warm latency histogram in pIC). **d**, Example traces (average of 5 trials) of transient (T) and sustained (S) responses to 10°C cooling (top) and warming (bottom); histograms of duration index for cooling and warming stimuli. Arrow heads indicate the example neurons. **e** Duration index of cool (top) and warm (bottom) responses plotted against the response latency. Green line at duration index 0.5 highlights the separation between T and S neurons. **f**, Duration index of warm versus that for cool in broadly tuned neurons (*n* = 125 cells). **g**, Left, example traces of cool (T and S) and warm (S) neurons responding to ∼10 s stimulus at fast onset speed (left, ∼130 C/s) or at slower rate (right, ∼1 C/s) (*n* = 5 trials). Below, corresponding stimulus traces. Right, histograms of adaptation index for T and S neurons separated accordingly to 0.5 duration index as in **e**. Arrow heads indicate example neurons. **h**, Model of how different channels of thermal afferent input could drive cortical neuron dynamics.

The spatial distribution of cortical neurons with respect to their functional properties is an important feature of cortical information processing (Chen et al., 2021, 2011; Ohki et al., 2005). To address whether pIC was thermotopically organized, we analyzed the spatial distribution of thermally tuned neurons in pIC (Fig. 2f, g, h). Visual inspection showed a heterogeneous arrangement of thermally responsive neurons (Fig. 2f). In agreement, the distance between similarly tuned neurons (cool to cool; warm to warm), were not significantly different to the distance between differently tuned neurons (cool to warm) (Fig. 2g). In a subset of experiments, we also used tactile stimulation and, in line with the differing spatial arrangements of touch and temperature, we observed a smaller fraction of thermally responsive pIC neurons (3%) than S1 neurons (12%) that responded to both thermal and tactile stimuli (Supplementary Fig. 3). Thus, thermal zones of pIC contain a heterogeneous, salt-and-pepper like, arrangement of thermally responsive neurons.

### Distinct temporal dynamics of cortical thermal responses

The skin contains discreet spots that evoke cool or warm sensations (Blix, 1882; Filingeri, 2016; Vriens et al., 2014). Interestingly, in vivo recordings from sensory afferent neurons that innervate the skin spots have shown distinct response dynamics to thermal stimuli with short latency, transient cool response mediated by Aδ-fibers, and longer latency, sustained firing to cool and warm stimuli by C-fibers (Darian-Smith et al., 1973, 1979; Duclaux and Kenshalo DR, 1980; Kenshalo and Duclaux, 1977). We went on to investigate whether distinct temporal dynamics between cool and warm were also present in cortical responses (Fig. 3a). In remarkable similarity to the periphery, a grand average of all significant cool and warm responses in pIC showed a shorter latency (∼80 ms) and more transient cool response compared to a longer latency (∼320 ms) and more sustained warm response (Fig. 3b). The latency difference is present in both tuned and broadly tuned neurons (Fig. 3c), suggesting that thermal responses are driven by similar input. Moreover, the cool response latency in S1 is similar to pIC, whereas the sparse warm responses in S1 are substantially delayed and variable compared to pIC warm responses (Fig. 2c; Supplementary Fig. 3).

We quantified the temporal dynamics of the thermal responses in pIC by computing a duration index which measures the change in response level at the end of the stimulation period compared to the initial peak value (Fig. 3d). The bimodality of the distribution of duration index for cool responses suggests a transient and a sustained response type (Fig. 3d). In contrast, warm responses show a broad distribution of sustained responses (Fig. 3d). Plotting the duration index against the response latency showed that the majority of cool transient neurons have a short latency onset, whereas both cool and warm sustained neurons show delayed onsets (Fig. 3e). In broadly tuned neurons, we observed a similar distribution of cool and warm dynamics with cool transient and warm sustained responses in the same cells (Fig. 3f), together highlighting that response dynamics are governed by sensory input rather than intrinsic cellular properties.

Fast onset, transient neuronal responses are thought to be reliable indicators of stimulus change, whereas sustained responses are optimal for absolute stimulus level encoding. We tested this hypothesis by measuring the decrease in response amplitude when a thermal stimulus is presented with different onset rates (adaption index in Fig. 3g, Supplementary Fig. 4). We observed that cool transient neurons were not activated by stimuli with a slow onset whereas sustained neurons responded similarly irrespective of the stimulus onset speed. These features were observed in highly tuned and broadly tuned cells (Supplementary Fig. 4). Overall, these data support a classic model of thermal encoding (Filingeri, 2016; Vriens et al., 2014), whereby cool responses are driven by a combination of fast Aδ-fiber and slow C-fiber afferent input and warm by slower C-fiber input (Darian-Smith et al., 1973, 1979; Duclaux and Kenshalo DR, 1980; Kenshalo and Duclaux, 1977; Paricio-Montesinos et al., 2020) (Fig. 3h).

### Different schemes for the encoding of cooling and warming amplitude

The amplitude of a sensory stimulus can be encoded in different ways: (i) by neurons with specific preferred amplitude values, (ii) the recruiting of more neurons as amplitude increases, (iii) graded rate coding (Fig. 4a). To address this in the thermal system, we measured single cell responses to different amplitude cooling and warming stimuli (Fig. 4b) and show that responses to different amplitudes were reliable within a neuron, but diverse between neurons (Fig. 4c). Plotting the population response amplitude showed an asymmetric response profile with a steeper curve for cooling and more graded for warming (Fig. 4d). The majority of cool neurons respond to the smallest amplitude tested (2°C) with only a minority recruited at larger amplitudes, whereas the number of activated warming neurons more gradually follows the increase of stimulus amplitude (Fig. 4e). The proportion of cool neurons reaching the maximal response amplitude is graded with stimulus amplitude, whereas warm neurons only reach their maximum response amplitude with large amplitude stimuli (Fig. 4f). Overall, these analyses suggest that the encoding scheme for thermal amplitude is a combination of extra recruitment of cells, model (ii), and graded changes of response amplitude, model (iii). Among the cells that responded significantly to warm, ∼5% showed an unexpected response profile with thermometer-like properties, responding to small but not large amplitude cooling. These ‘thermometer cells’ showed graded reporting of absolute thermal value, independent of the stimulus direction from the AT (Fig. 4c, cell #5; Supplementary Fig. 5).

**Figure 4.**
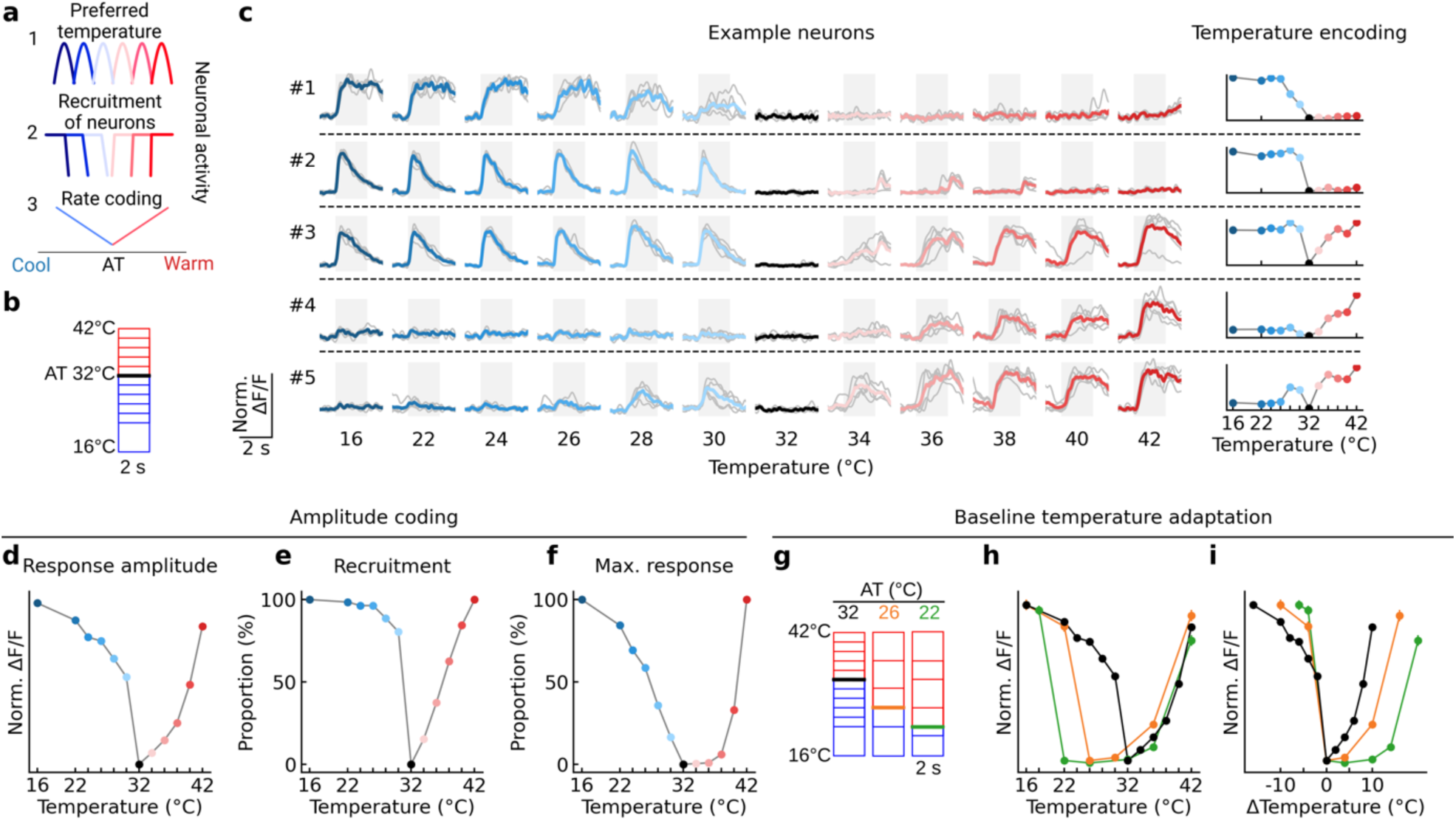
Relative vs. absolute encoding of cooling and warming. **a**, Schematic showing three possible cortical thermal coding schemes. **b**, Schematic of stimulus protocol from AT of 32°C. **c**, Left, traces of example neurons responding to cooling and warming from AT 32°C, grey lines shows individual trials (5 trials per temperature), colored shows mean response. Right, peak response amplitude plotted as a function of the thermal stimulus for the example neurons on the left. **d**, Summary of response amplitude plotted as a function of the thermal stimulus for entire population of cells (*n* = 746 cells, 16 sessions, 7 mice). Colored filled circles show mean ± s.e.m. at AT 32°C. **e**, Same data as **d**, but showing the proportion of recruited neurons (> 20% of response amplitude). **f**, Same data as **d**, but showing the proportion of neurons reaching max response (> 80% of response amplitude). **g**, Schematic of stimulus protocols used to test impact of AT on thermal encoding. **h**, Left, graphs as in **d**, but for all ATs studied. (T 32°C *n* = same as **d**; AT 26°C *n* = 401 cells, 9 sessions, 5 mice; AT 22°C *n* = 448 cells, 10 sessions, 6 mice). **i**, Same data as **h** but plotted against change in stimulus amplitude.

Primary sensory afferents and spinal cord neurons thermal responses are influenced by the skin AT (Andrew and Craig, 2001; Craig et al., 2001; Darian-Smith et al., 1973, 1979; Duclaux and Kenshalo DR, 1980; Kenshalo and Duclaux, 1977; Ran et al., 2016; Wang et al., 2018; Yarmolinsky et al., 2016), but whether this is reflected by a change in cortical response amplitude it is not known. To investigate this, we compared responses to the same amplitude stimuli but from ATs of 22 and 26°C (Fig. 4g). At lower ATs, warming stimulus amplitude had to be substantially increased to observe response amplitudes comparable to those seen at AT 32°C and responses were often only observed once the stimulus target temperature reached 34°C. In contrast, reliable responses to cooling were observed for all target temperatures and ATs tested (Fig. 4h). This difference was highlighted by plotting the response amplitude against the relative stimulus amplitude which showed a shift in the warm response curve, but not in the cool (Fig. 4i, consistent conclusions were also observed in widefield imaging data in Supplementary Fig. 6). Together these data indicate that cooling and warming amplitude is encoded in fundamentally different ways in the cortex, with warming dependent on the absolute temperature and cooling reflecting the magnitude of temperature change, consistent with findings in primary sensory afferents and spinal cord neurons (Andrew and Craig, 2001; Craig et al., 2001; Darian-Smith et al., 1973, 1979; Duclaux and Kenshalo DR, 1980; Kenshalo and Duclaux, 1977; Ran et al., 2016; Wang et al., 2018; Yarmolinsky et al., 2016).

### Thermal perception is mediated by posterior insular cortex

More anterior parts of insular cortex are thought to play a role in cognitive and motivational control of behavior (Craig, 2002; Gogolla, 2017), while a central region is involve in taste perception (Peng et al., 2015). Human lesions, microstimulation and imaging studies have suggested a role for insular cortex in thermal perception (Birklein et al., 2005; Craig et al., 2000; Penfield and Faulk, 1955), but reversible manipulations have not been performed and the link between thermal sensitivity and perception is unclear. To address this, we performed optogenetic manipulations during a Go/NoGo thermal detection task (Fig. 5a). Thermal stimuli were delivered at random times from AT 32°C. VGAT-ChR2 mice, with channelrhodopsin-2 (ChR2) constitutively expressed in GABA-ergic inhibitory interneurons, were implanted with an optical window over pIC and provided with water rewards upon licking a water spout during the onset ramp or plateau phase of the stimulus (Fig. 5a, Supplementary Fig. 7). Because of differences in sensitivity to cooling and warming in pIC (Fig. 4), we chose 10°C, 2°C, 1°C and 0.5°C cooling stimuli, whereas for warming we used 10°C, 8°C, 6°C, and 4°C. Mice were able to report stimuli at all amplitudes highlighting their acute thermal perceptual ability (Fig. 5b and c). Similar to the functional responses (Fig. 1 and 3), mice reported warming at longer latencies than for cooling. We went on to inhibit the activity of pIC during the stimulus phase in the same mice (Fig. 5a, e and f) using light pulses (12.5 mW, 20 Hz, 2 s) via a 200 µm fiber optic positioned normal to the thermal region of pIC (Fig. 5a, e and f). In cooling trained mice, optogenetic inhibition of pIC suppressed the hit rates to 2°C, 1°C and 0.5°C stimuli but not to 10°C (Fig. 5e); whereas, in warming trained mice, optogenetic inhibition suppressed the hit rates to all amplitudes (Fig. 5f). The different impact of optogenetic inhibition to warming and cooling possibly result from the additional representation of cool in S1 or the fast onset of cool responses allowing more robust encoding for cool than warm. Repeating the same optical stimulation paradigm but in Thy1-GCaMP6s mice not expressing ChR2 showed no effect on thermal perception (bottom panels Fig. 5e, f; (Supplementary Fig. 7), confirming that the light alone does not alter the behavior of the mice in our task. Moreover, optogenetic inhibition of pIC in VGAT-ChR2 mice during spontaneous licking of free water rewards and during an acoustic detection task did not alter lick rates (Supplementary Fig. 7). Finally, optogenetic inhibition of pIC in VGAT-ChR2 mice trained to discriminate between and acoustic stimulus (Go; rewarded) vs. an acoustic stimulus presented simultaneously with a thermal stimulus (NoGo; not rewarded) induced a selective increase of licking during the NoGo trials, showing that pIC manipulation has an impact on perception rather than a general decrease of licking (Supplementary Fig. 7). Overall, our data show that pIC plays a profound causal role in thermal perception.

**Figure 5.**
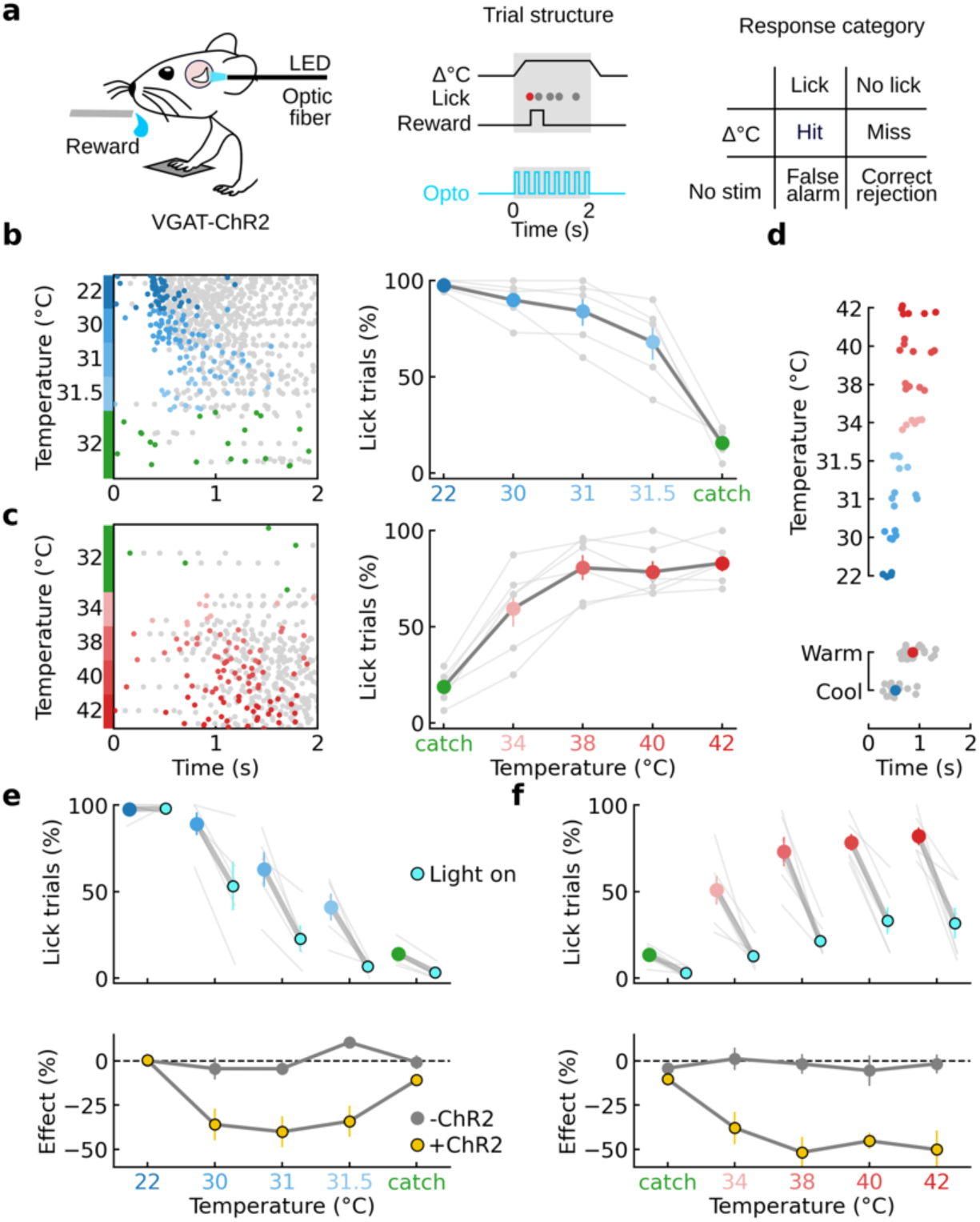
Thermal perception is mediated by posterior insular cortex. **a**, Left, schematic of detection task and placement of the optic fiber. Middle, example trial structure showing timing of reward window (gray) and the timing of an optical stimulus during trials with optogenetic manipulation. Filled circles show licks, first lick colored to show rewarded lick. Right, response categories of task. **b**, Left, raster plot of licks in all trials (*n* = 300) from an example mouse trained for cooling sorted by stimulus amplitude, green filled circles show false alarms. Right, summary of behavioral performance showing proportion of trials with at least one lick in reward window (*n* = 5 mice). **c**, Same as **b**, but for warming (raster *n* = 239 trials, graph *n* = 6 mice). **d**, Latency of first lick is longer to warming than cooling at different amplitudes. Filled circles show data from individual mice in **b, c**. Bottom, data from all amplitudes with mean ± s.e.m. **e**, Top, proportion of trials with licks in VGAT-ChR2 mice (*n* = 5) during optical stimulation (cyan) trials vs. without optical stimulation (blue) for different cool stimulus amplitudes. Gray lines show individual mice, colored filled circles show mean ± s.e.m. Bottom, shows the effect of light stimulus (change in percentage of trials with licks, mean ± s.e.m.) from the VGAT-ChR2 mouse (yellow), and from mice not expressing ChR2 (gray, see Supplementary Fig. 7). **f**, Same as **e**, but for warm stimulation (*n* = 6 mice).

## Discussion

Here we identify a cortical region required for non-painful thermal perception and support the hypothesis that pIC houses a thermal cortex. Our data show that humans, monkeys and rodents therefore not only share similar perceptual abilities (C. Stevens Kenneth K. Choo, 1998; Paricio-Montesinos et al., 2020; Rózsa et al., 1985), but also a cortical area (pIC) specialized in temperature processing, putting the thermosensory system of mammals closer than previously thought (Craig, 2009; Craig et al., 2000). Our data describe distinct encoding features of cool and warm in the cortex. The representation of warm has delayed and uniform dynamics that encodes absolute stimulus amplitude, compared to the mixed temporal dynamics for cool that drive a relative encoding of stimulus amplitude. Similar features have been reported in primate and human primary thermal afferent neurons (Darian-Smith et al., 1973, 1979; Duclaux and Kenshalo DR, 1980; Kenshalo and Duclaux, 1977) and closely resemble thermal response features in *Drosophila* central and peripheral neurons (Frank et al., 2015; Liu et al., 2015), together highlighting the remarkably conserved nature of thermal encoding across the animal kingdom.

We find that cool and warm are cortically represented largely following labelled line principles (Fig. 1a - top)(Andrew and Craig, 2001; Craig et al., 2001; Darian-Smith et al., 1973, 1979; Duclaux and Kenshalo DR, 1980; Kenshalo and Duclaux, 1977; Ran et al., 2016; Wang et al., 2018; Yarmolinsky et al., 2016). The distinct features of cool and warm in broadly tuned neurons show that functionally segregated streams of afferent information can merge in cortical neurons while conserving the features of tuned neurons. This is interesting in light of recent data indicating that tactile input undergoes substantial subcortical transformation before reaching S1 (Emanuel et al., 2021). Nevertheless, cortical neurons can show responses features like the ‘thermometer cells’ not observed in the periphery or in highly tuned cortical neurons, raising the possibility of cortical generation of complex thermal features (Fig. 4c, Supplementary Fig. 5).

Touch and cool response areas overlap in S1 (Milenkovic et al., 2014), whereas they are separate in pIC, hinting at functional differences. Objects are normally cooler than skin temperature and thermally conductivity is an important component of object identification (Ho and Jones, 2006). The cool-only representation in S1 could therefore be to integrate cool with touch during haptic sensing, whereas pIC could encode the temperate level. Taken together, the discovery of an optically accessible cortical representation of temperature, provides a platform to address the neural mechanisms non-painful thermal perception, as well as pain evoked during noxious stimulation, allodynia and thermal illusions.

## Acknowledgements

This work was supported by the European Research Council (ERC-2015-CoG-682422, J.F.A.P.), the European Union (3×3Dimaging 323945, J.F.A.P.), the Deutsche Forschungsgemeinschaft (DFG, FOR 2143, J.F.A.P., SFB 1315, J.F.A.P.), the Helmholtz Society (J.F.A.P.), the Centre National de la Recherche Scientifique (CNRS) (M.C.) and the Independent Research Fund Denmark (M.V.). We thank Michael Brecht, Gary Lewin, Jens Kremkow, Sylvain Crochet and members of the Poulet lab for constructive comments on an earlier version of the manuscript, Svenja Steinfelder for help with administrative and technical aspects, Janett König for help with immunohistochemistry and Prof. Dr. Britta Eickholt for sharing NeuN antibody.

## Author contributions

M.V., M.C. and J.F.A.P. designed the study. M.V. and M.C. performed experiments and analyzed the data. M.V., M.C. and J.F.A.P. wrote the manuscript. The authors declare no competing financial interests.

