## Supplementary Information for "The cellular coding of temperature in the mammalian cortex"

M. Vestergaard, M. Carta and J.F.A. Poulet

**Supplementary information contains:**

Methods

Supplementary Figs 1 – 7

Supplementary References

### Methods

#### Mice

All experimental procedures were carried out in accordance with the State of Berlin Animal Welfare requirements and European animal welfare law. Male and female mice older than 2 months were maintained on a 12:12 hr light-dark cycle with experiments performed during the light phase of the cycle. Mice were housed in groups with *ad libitum* access to food and water unless stated. Thy1-GCaMP6s mice (C57BL/6J-Tg(Thy1-GCaMP6s)GP4.3Dkim/J) were used for calcium imaging and for behavioral experiments (The Jackson Laboratory, stock no. 024275) (Chen et al., 2013; Dana et al., 2014). VGAT-CHR2 (B6.Cg-Tg(Slc32a1-COP4*H134R/EYFP)8Gfng/J were used for behavioral experiments (The Jackson Laboratory, stock no. 014548) (Zhao et al., 2011). For awake experiments, mice were gradually habituated to head- and paw-fixation and tilted for optical access to pIC or S1.

#### Surgery

Mice were deeply anesthetized (ketamine 120 mg/kg and xylazine 10 mg/kg) and injected subcutaneously with dexamethasone sodium phosphate (2 mg/kg) to prevent cerebral edema, as well as metamizol (200 mg/kg). Anaesthetized mice were fixed with a nose-clamp, eye gel (Vidisic, Bausch + Lomb) applied to both eyes and body temperature maintained at 37°C with a heating pad and rectal probe. After surgery, mice received a subcutaneous injection of warm sterile saline solution and were placed on a warm blanket until they awoke from anesthesia. To avoid post-operative pain, metamizole was dissolved in the drinking water for 3 days post-surgery.

*Implantation of cranial windows*

To implant a window over posterior insular cortex (pIC), mice were anaesthetized as above, then the parietal and temporal bones of the left hemisphere were exposed and the left temporalis muscle was carefully separated from the temporal bone and partially removed. A ~3x3 mm craniotomy was performed over pIC as identified by local anatomical landmarks (rhinal vein, middle cerebral artery, zygomatic bone). To implant a window over primary somatosensory forepaw cortex (S1) a 3 mm diameter craniotomy was performed over the left hemisphere S1 identified by the anatomical location respect to bregma (1.5-2.5 mm lateral and 0.5-1 mm anterior). The cortical surfaces were then rinsed with Ringer's solution and 2 glass coverslips were placed on the surface of the cortex. The lower glass (diameter 3 mm or 3x3 mm) and the upper glass (diameter 4 or 5 mm) were glued together with optical adhesive (NOA 61, Norland Products). Dental cement and cyanoacrylate glue were used to attach a metal head post and the upper coverslip to the skull. Following pIC surgery, the skin lying over the temporal muscle was sutured.

*Clear skull preparation*

To gain simultaneous optical access to S1 and pIC, we developed a through-skull preparation. Under deep isoflurane anesthesia, the left temporalis muscle was partially removed and the dorsal surface of the skull was cleared of skin and periosteum. Finally, the exposed skull was sealed with cyanoacrylate glue (Fig. 1f, g).

**Cortical pharmacology**

For pharmacological inactivation ~500 µm craniotomies were drilled over cool (32-22°C) responsive areas in S1 and pIC in mice anesthetized with isoflurane (1.5-2% in $O_2$). Both regions were identified functionally using the widefield calcium imaging response to thermal stimuli. Following the craniotomy, the dura was covered with transparent silicone gel (3-4680, Dow Corning, Midland, MI). 300 nl of muscimol or Ringer's solution were injected at a rate of 100 nl/min 300-500 µm below the pial surface using a pulled glass pipette and a hydraulic injection system (MO-10, Narashige). Muscimol (Abcam, ab120094) was dissolved in Ringer's solution to a concentration of 5 mM. Imaging sessions were performed > 10 minutes followed the end of Ringer's solution or muscimol injection (Fig. 1f, g).

**Sensory stimulation**

*Thermal*

For widefield imaging and behavioral experiments, thermal stimuli were delivered by an 8x8 mm Peltier element regulated by a feedback-controlled stimulator (Yale Medical School or a custom-made device, ESYS GmbH Berlin). Different thermal stimuli (2 s duration with an onset ramp speed of 20°C/s) were interleaved randomly and delivered every 30 or 60 s. During paw stimulation, we took care to place the fore- or hindpaw pad glabrous skin into direct contact with the center of the Peltier element. During face stimulation, the Peltier surface was positioned on the orofacial region of the snout. For single cell two-photon experiments thermal stimuli were delivered using a QST-lab thermal stimulator (https://www.qst-lab.eu/). Thermal stimuli of different durations (2, 5, 10 s) or with different onset ramp speeds (approximately 130, 10, 3.3, 2, 1 °C/s) were interleaved randomly and delivered every 30 or 60 s.

*Acoustic*

8 kHz, ~65 dB SPL, 1 s duration acoustic stimuli were delivered every 60 s using a loud speaker (Visaton) positioned 10 cm away from the contralateral ear. 8 kHz was chosen because is well represented in the pIC auditory field (Sawatari et al., 2011). For behavioral training, a 14 kHz tone (~65 dB, 1 s duration) was chosen as it is weakly represented in the pIC auditory field (Sawatari et al., 2011).

*Vibrotactile*

Vibrotactile stimuli (100 Hz) were generated by a piezoelectric actuator (PL127, PI) equipped with a 5 cm glass rod bent at the tip. The contralateral fore- or hindpaw was tethered to a rigid support with a hole (diameter 5 mm) to allow the bent tip of the glass rod to stimulate the center of the palm of the paw. In some experiments vibrotactile stimuli were generated using a small motor (Pololu shaftless vibration motor 8x3.4 mm) positioned in top of the paw tethered to the Peltier. Vibrotactile stimuli had a 0.5 s duration and were delivered every 30-60 s.

**Widefield imaging**

*Setup*

Imaging was performed with a Leica MZ10F stereomicroscope. Blue excitation light (470 nm) was emitted from a LED (pE-300, CoolLED) and band pass filtered (470/40 nm). Emission light was band pass filtered (525/50 nm) before recording with a sCMOS camera (ORCA-Flash4.O LT, Hamamatsu). Images were acquired at a rate of 20 Hz and 35 ms exposure time. Frame size was 1024x1024 pixels (2x2 binning) and for cranial window preparation the field-of-view was ~2.7x2.7 mm and for the cleared skull preparation the field-of-view was ~6.7x6.7 mm. Acquisition trials lasted between 10-15 s and had an inter-trial-interval time of 30-60 s. During pIC imaging, animals were tilted on their right side to allow optical access. Data acquisition was controlled by custom written code (Python).

*Widefield image analysis*

Movies were motion corrected and areas in the recordings outside the brain were masked. We calculated the relative change in fluorescence as $\Delta F/F = (F(t) - F_0)/F_0$ where $F_0$ is the 15$^{th}$ percentile of the activity in the trial. The average activity of the last 500 ms before stimulus were subtracted from ΔF/F. The activity of a given region was quantified by averaging the fluorescence in a region of interest (ROI, diameter 15 pixels) before calculating the ΔF/F. When placing ROIs we avoided visible blood vessels, but otherwise positioned the ROI at the area with the peak stimulus driven response. The ROI positions were kept the same over days/recording sessions by comparing the position relative to the blood vessels. Auditory stimuli activated several fields in the auditory cortex (AC) and one area in pIC (insular auditory field, IAF).

*Somatotopic maps*

Somatotopic maps of sensory input to pIC in Fig. 1 are grand averages across mice and trials. Not all stimuli were delivered to all mice. Using the ROI of the thermal forepaw response as a reference location, ΔF/F movies for all trials were aligned by translation. The grand averages were smoothed with Gaussian filter ($\sigma$ = 20 pixels) and threshold to only show peak activity (> 85% of the max activity for thermal and > 90% for tactile stimulation). Because of the simultaneous activity in the IAF and AC with sound stimuli their responsive responses were analyzed independently.

*Response onset estimation*

We defined the onset of the widefield response as the peak of the second derivative of the widefield signal in a time window from the start of stimulus to the peak of the first derivative. If negative values of the first derivative appeared after stimulus start, the time window was from the last of such negative values. We only performed this analysis on data with a response amplitude > 3% ΔF/F.

**Two-photon imaging**

*Setup*

Fluorescence signals were recorded using a two-photon microscope (ThorLabs Bergamo II, 12 kHz scanner) with a Nikon 16x water-immersion objective (NA 0.8) giving a field-of-view of 540x540 µm. The scope was operated using ThorImageLS software (Thorlabs). In the majority of the experiments the microscope was rotated by 45° from vertical. The excitation laser (InSight DeepSee, Spectra-Physics) was tuned to 940 nm and the power never exceeded 100 mW. Emitted photons were bandpass filtered 525/50 (green) onto a GaAsP photomultiplier tube. Multi-plane (512x512 pixels) acquisition was controlled by a fast piezoelectric objective scanner, with planes spaced 45 µm apart in depth. 7 planes were acquired sequentially, and the scanning of the entire stack was repeated at ~5 Hz. Beam turnarounds at the edges of the image were blanked. Acquisition trials lasted 26 s and had an inter-trial interval of 30-60 s. Stimuli generation and hardware synchronization were performed on a computer with a National Instrument card running custom written Python code.

*Two-photon analysis*

Motion correction of data, identification of putative neurons and calculation of ΔF/F was performed using the Suite2p package in Python (Pachitariu et al., 2016). Identified neurons were manually verified by visually examination of the traces and the spatial footprints of the neurons. A Savitzky–Golay filter was applied to traces presented in figures for visual purpose only.

*Criteria for responsive neurons*

Neurons with a significant increase in activity to 10°C cooling and warming stimuli were included for further analysis. Cool and warm neurons were defined as those significantly responding to either to 10°C cooling or warming stimuli. Broadly tuned neurons had significant response to both cooling and warming stimuli. To identify an increase in response amplitude, the activity in a time window before stimulus onset was compared with the activity in a time window during stimulus. A neuron was identified as being responsive if the change in activity was significantly larger (estimated by bootstrapping) than 2 times the noise level of the neuron measured by the interquartile range of activity before the stimuli.

The Thermal bias index (Fig. 2d, e) for a thermally responding neuron is the normalized difference between its cool and warm responses, (warm – cool) / (warm + cool), where "warm" is the average peak response to 32-42°C and 'cool' is the average peak response to 32-22°C. The Duration index (Fig. 3d) measures the change in the fluorescence level between the initial peak value ($f_{init}$) and the end ($f_{end}$) of a 2 s thermal stimulus, ($f_{init}$ – $f_{end}$) / $f_{init}$. The Adaptation index (Fig. 3g) was calculated as the difference in the max. fluorescence level during fast thermal stimulation (~130 °C/s)($f_{fast}$) and during slow thermal stimulation (~1 °C/s)($f_{slow}$), ($f_{fast}$ – $f_{slow}$) / $f_{fast}$.

**Behavioral experiments**

*Setup*

Head fixed mice (VGAT-CHR2 and Thy1-GCaMP6s) were implanted with a glass window over pIC and trained on a go-nogo stimulus detection paradigm. Mice were rewarded with a water drop (~2 µl) if they reported a randomly timed thermal stimulus with at least one lick of a water spout (capacitance sensor) within a window of opportunity (from the start of thermal stimulus to start of offset ramp). Correct rejections were not rewarded and incorrect responses were not punished, although premature licking in the 5 s before the stimulus onset would postpone the next trial by 5 s. Mice were water restricted and their weight monitored daily. For behavioral training, a 200 µm diameter, 0.22 NA optic fibre (Thorlabs) was coupled to an LED (470 Plexon LED source) and placed on the coverslip surface orthogonal to the thermal region of the pIC using the blood vessel pattern. Control of behavioral training and data collection was performed using custom-written Labview software (National Instruments, USA).

*Thermal perception task*

Thermal perception (Fig. 5) training involved several stages: (1) Free access to water rewards from the water spout. (2) Automatic water rewards paired to the presentation of 10°C, 2 s cooling and warming stimuli from adapting temperature (AT) 32°C. (3) Training to report a 10°C cooling and warming stimuli by licking for a water reward within a 2 s window from the start to the end of the plateau phase of the stimulus. Thermal stimuli trials and catch trials were presented with a ratio of 1:1 with an inter-stimulus-interval of 15-20 s. (4) Once a hit rate > 70% and false alarm rate < 30% had been reached, 4 randomized stimulus amplitudes were used from AT 32°C (cooling, 22°C, 30°C, 31°C and 31.5°C; warming 42°C, 40°C, 38°C and 34°C) with a 4:2 ratio of stimulus trials to catch trials. (5) Once a hit rate of > 70% and false alarm rate < 30% had been achieved for at least 1 amplitude, in the next session we presented the same 4 thermal stimuli with 1 catch trial with LED on or off (ON/OFF trials) with a ratio of 2:1 (randomized). The LED was on for the entire duration of the stimulus and delivered at 20 Hz, 50% duty cycle with a power of 12.5 mW (measured at tip of fiber). Prior to the start of stage (4) or (5) sessions, mice were exposed to a brief session of the stage (3) protocol to quench initial thirst.

*Thermal discrimination task*

Discrimination task (Supplementary Fig. 7) training involved several stages: (1) Free access to water rewards from the water spout. (2) Automatic water rewards paired to the presentation of 14 kHz, ~65 dB, 0.5 s acoustic stimulus. (3) Training to report 14 kHz stimulus by licking for a water reward within a 1.4 s window from the start of the stimulus (Go trial). Acoustic stimuli (14 kHz, ~65 dB) delivered 0.6 s after the beginning of a 2 s thermal stimulus (either cooling or warming) (NoGo trial) and catch trials (no stimuli) were not rewarded. Go, NoGo and catch trials were presented with a ratio of 1:1:1 with an inter-stimulus-interval of 15-20 s. Training was continued until Go trial hit rate > 70%, NoGo false alarm rate around 50% and false alarm catch rate < 30%. (4) Once a mouse learned to discriminate between acoustic stimulus alone (Go) and acoustic stimulus presented together with thermal stimuli (NoGo), in the next session we presented the same stimuli interleaved at a ratio of 2:1 into trials with the LED off (OFF) and trials with the LED on (ON). The LED was on for the entire duration of the thermal stimulus and delivered at 20 Hz, 50% duty cycle with a power of 12.5 mW (measured at tip of fiber). Prior to the start of stage (4) sessions, mice were exposed to a brief session of the stage (3) protocol to quench initial thirst.

*Acoustic perception*

Acoustic training with optogenetic manipulation had the same structure as the thermal training above, but with a 14 kHz, ~65 dB, 2 s acoustic stimulus.

*Free licking*

To monitor the impact of pIC optogenetic manipulation on licking behavior, water restricted mice were allowed to freely lick from a lick spout with continuous rewards. Licking rates during 30 s trials with no LED stimulation were compared to trials with 5 s, 12.5 mW, 20 Hz, 50% duty cycle LED stimulation trials.

*Data sets and analysis*

Mice were randomly allocated to be trained first to report cooling or warming. In a second training round, they were trained to report the opposite stimulus. Data measuring the thermal perceptual ability of mice without optogenetic manipulation (Fig. 5b, c, d) is generated from the same mice used in optogenetic manipulations (Fig. 5 e,f), the day before optogenetic testing. The effect of the optogenetic manipulation (bottom panels of Fig. 5e, f and Supplementary Fig. 7) was quantified by the mean change in the percentage of lick trials.

**Histology**

In order to mark the thermal area of cortex, animals were deeply anesthetized by intraperitoneal injection of ketamine/xylazine and the cranial window gently removed. Next, the tip of a fine glass pipette painted with fluorescent dye (DiI, 5 mg/ml) was inserted into the center of the functional response area and left in place for 5-10 minutes. Mice were then perfused transcardially using cold phosphate buffered saline (PBS, 0.1 M, pH 7.4) and fixed with paraformaldehyde (PFA, 2% in 0.1 M PBS). The brain was removed and kept for 1-5 hours in 2% PFA. After washing with PBS, the hemispheres were separated and cortices were removed and flattened between 2 glass slides separated by a spacer of 1-2 mm. Glass slides were weighed down for approximately 3-8 hrs at 4°C in 2% PFA. After washing with PBS 70-80 μm sections were cut on a Vibratome (Leica VT1000s). Sections were stained for cytochrome-oxidase activity (2 mg cytochrome C, 6 mg diaminobenzidine DAB). After the staining procedure, sections were mounted on glass slides with Mowiol mounting medium. Images were acquired with a Zeiss microscope (AX10) using a 5x objective and processed using Fiji (ImageJ, NIH, USA). Area boarders were manually delineated following the contrast of the cytochrome-oxidase stain and are comparable to previously reported data (Gămănuţ et al., 2018). 50 µm coronal sections of PFA-fixed mouse brains were stained for 48 hours with the following primary antibodies: mouse anti-Gad67 (cat. no.: MAB5406; Millipore; 1:800); chicken anti-GFP (cat. no.: ab13970; Abcam; 1:250); mouse anti-NeuN (cat. no.: clone A60 #MAB377; Millipore; 1:100; a gift from Britta Eickholt, Charite, Berlin). The secondary antibodies Cy3 goat anti mouse (A-21422; Invitrogen; 1:250) and A488 goat anti chicken (A-11039; Invitrogen; 1:250) were incubated for few hours at room temperature. Cell counting was performed manually in FIJI using the Cell Counter plug-in on epifluorescence images acquired using a Zeiss microscope (AX10) using a 10X objective.

#### Data analysis and statistics

No statistical methods were used to predetermine sample sizes. Experimenters were not blind to trial order, but trial order was randomized during the experiment. All data analysis was performed using Python. Uncertainty of means are reported with standard error of the mean. Bootstrapping was used for estimating central 95% confidence intervals using 5,000 resamples.

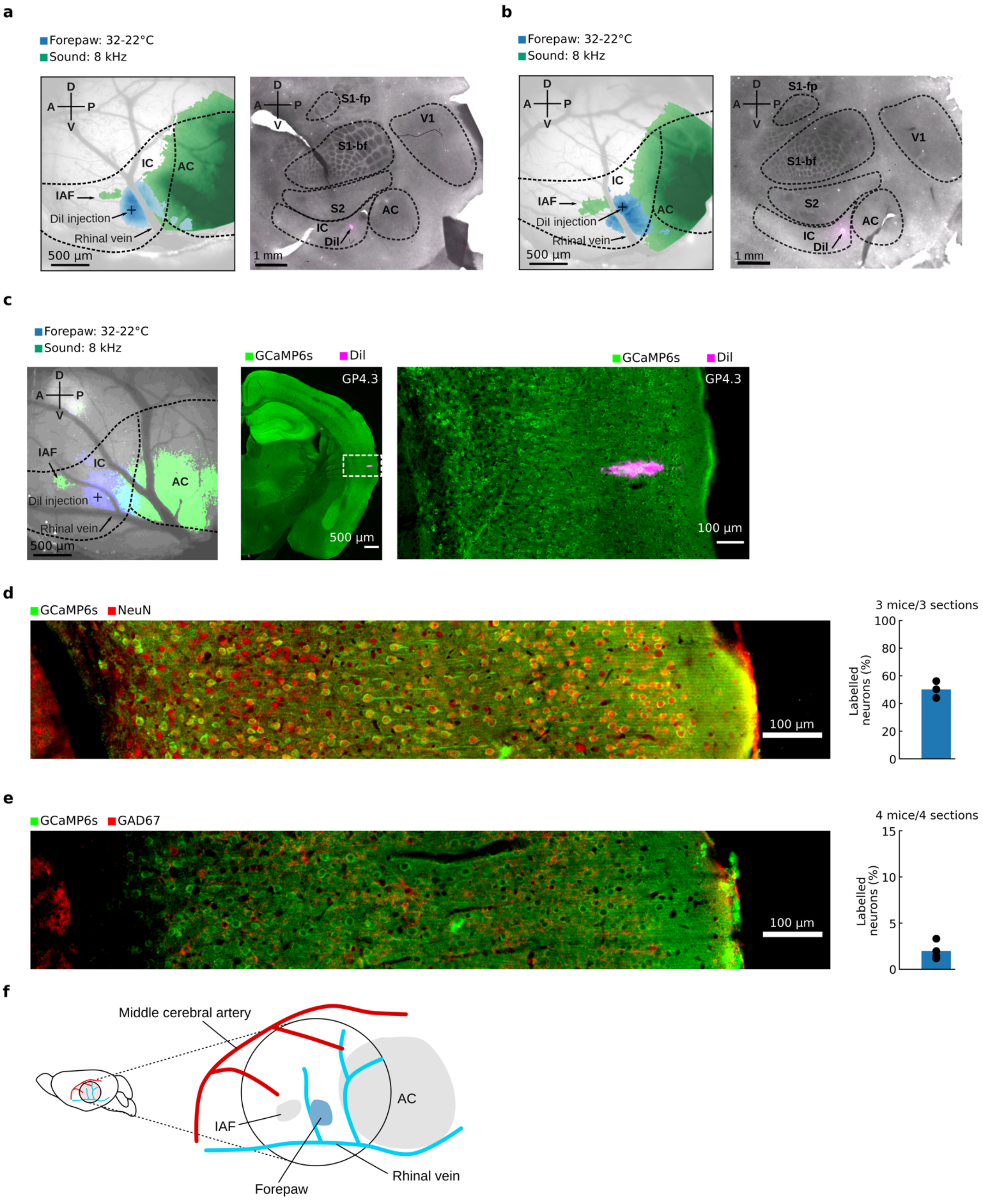


**Supplementary Figure 1. Functional and histological localization of pIC**

**a**, Functional and histological localization of the thermal zone of forepaw pIC in 2 example mice. Left, in vivo image of cortical surface of a Thy1-GCaMP6s mouse with superimposed colored areas showing the widefield responses to cool (blue, 32-22°C) and sound (green, 8 kHz). Glass pipette painted with Dil (5 mg/ml) was inserted in the center of the cool response area (black cross). Right, cytochrome oxidase staining of a flattened cortex preparation from the same mouse on left with site of Dil injection (magenta). Areal borders were manually delineated based on CO intensity reaction, parcellation followed *(Gămănuţ et al., 2018)*.
**b**, Second example mouse as in **a**.
**c**, Left, in vivo image of cortical surface of a Thy1-GCaMP6s mouse with superimposed colored areas showing the widefield responses to cool (blue, 32-22°C) and sound (green, 8 kHz). Center, coronal section of the same mouse on the left containing the site of Dil injection (magenta). GCaMP6s signal was amplified with immunohistochemistry with anti-GFP antibody. Right, zoom of the injection site. Example mouse is the same presented in Fig. 2c.
**d**, Left, example of epifluorescence image of pIC from a Thy1-GCaMP6s mouse stained for GCaMP6s (labelled with anti-GFP antibody) and for the general neuronal marker NeuN. Right, quantification of the percentage of NeuN-positive neurons also expressing GCaMP6s ($n$ = 3, mean). Values are in line with previous reports *(Dana et al., 2014)*.
**e**, Same as d but with staining for GCaMP6s and for the general GABAergic neurons marker GAD67. Right, percentage GAD67-positive neurons among GCaMP6s-expressing neurons ($n$ = 4 mice, mean). Values are in line with previous reports *(Makino et al., 2017)*.
**f**, Cartoon schematic representing the position of the acoustic and forepaw cool responsive fields relative to major blood vessels. IAF, insular auditory field; AC, auditory cortex.
Different terminology has been used for this region in prior studies, including pIC, parietal ventral area (PV) or S2/IC, highlighting the difficulties in discerning two neighboring cortical regions. In agreement with studies that have used functional mapping of sensory responses *(Gogolla, 2017; Rodgers et al., 2008; Sawatari et al., 2011; Zhang et al., 2020)* and because pIC is a region associated with thermal processing in humans studies *(Birklein et al., 2005; Bokiniec et al., 2018; ("Bud") Craig, 2018; Craig et al., 2000; Filingeri, 2016; Mazzola et al., 2012)* here we use the term pIC.

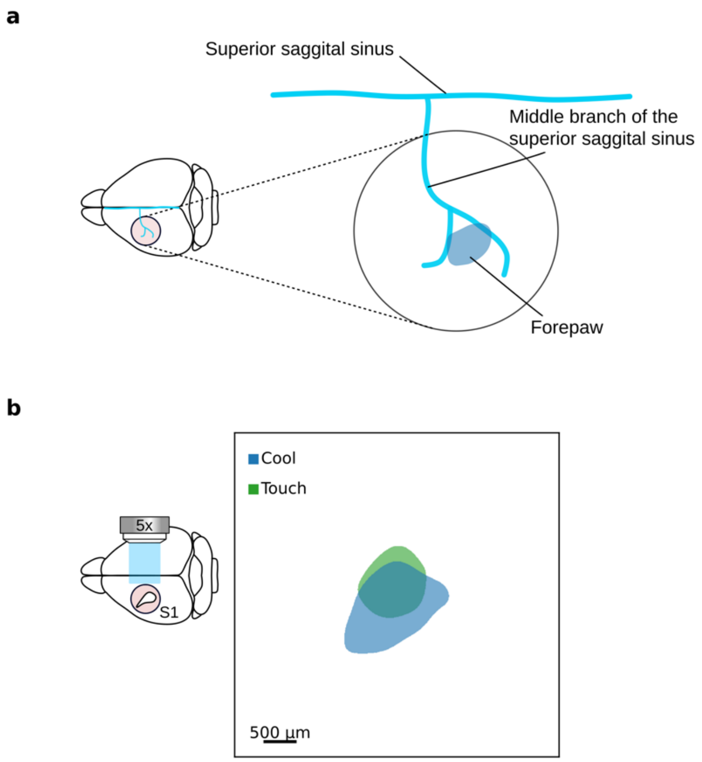


**Supplementary Figure 2. Overlap of tactile and cool representations in forepaw S1**
**a**, Cartoon schematic showing the location of the forepaw cool responsive area (blue) relative to the major blood vessels.
**b**, Cool (32-22°C) and tactile responses overlap in forepaw S1. Colored area indicates peak widefield calcium imaging response averaged across mice (*n* = 7 for cool and touch). Data from individual mice are aligned to peak activity of the cool forepaw response. Data confirm previous observations (Milenkovic et al., 2014).

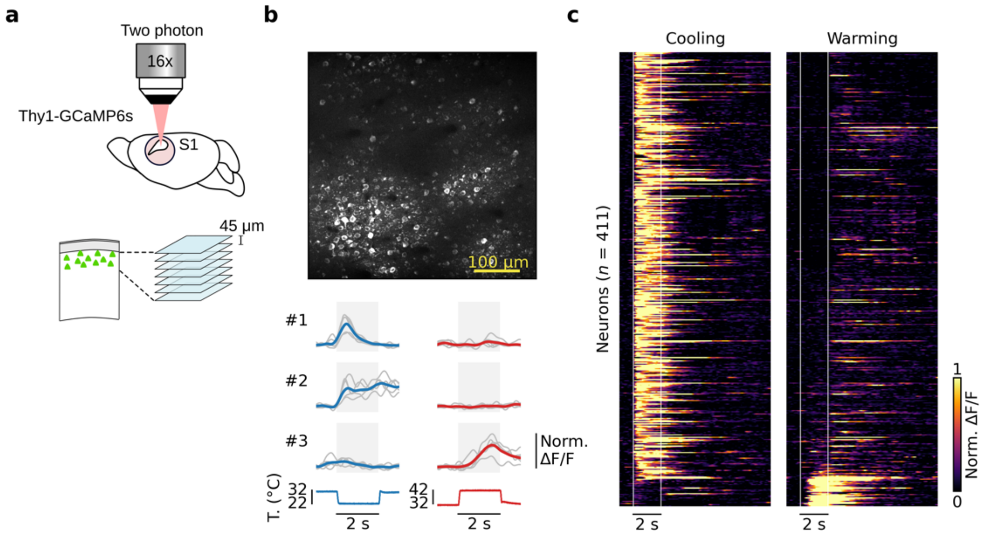


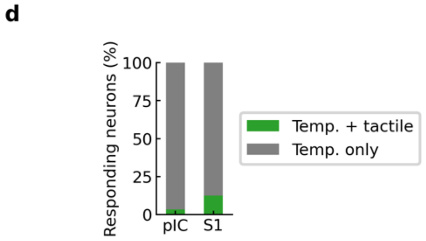


**Supplementary Figure 3. Cellular encoding of cooling in S1**

**a**, Cartoon schematic showing two-photon imaging of S1. Imaging started at 100 µm from the pial surface of a Thy1-GCaMP6s mouse, 7 optical sections were acquired with intervals of 45 µm.

**b**, Top, example in vivo two photon image of GCaMP6 expressing neurons in S1. Bottom, example of responses of single S1 neurons during 10°C cooling (blue) or 10°C warming (red) stimuli. Here and in all figures, gray lines show single trial responses, colored lines show average, grey area indicates time from start of stimulus to end of plateau phase. Below, corresponding stimulus traces.

**c**, Single cell S1 calcium responses to cooling (left) and warming (right) stimuli. Each line represents a single neuron, responses are normalized to the peak and sorted based on the thermal bias index with white lines showing the onset and the end of the plateau phase of thermal stimuli ($n$ = 411 cells, 4 mice, 9 sessions).

**d**, bar graph showing the percentage of pIC and S1 neurons responding only to thermal stimuli or also to tactile stimulation (pIC, 506 neurons thermal only, 18 neurons thermal and tactile; S1, 345 neurons thermal only, 49 neurons thermal and tactile).

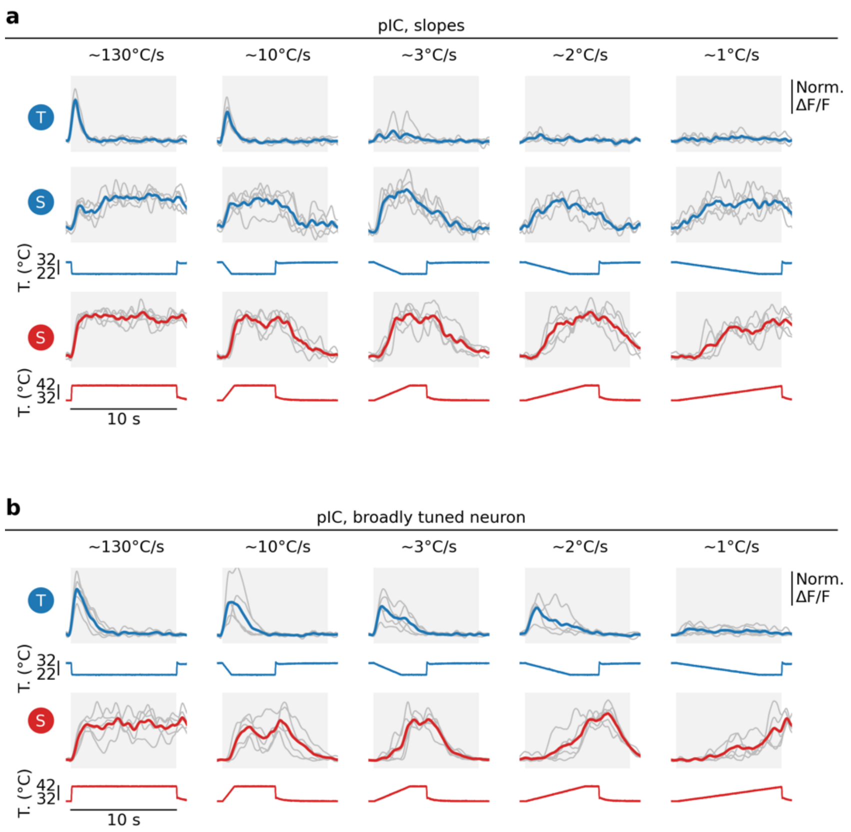


**Supplementary Figure 4. Example cells responding to thermal stimuli with different stimulus onset speeds**
**a**, Example traces of transient and sustained cool (T and S) and sustained warm (S) neurons responding to stimuli with different onset speeds of approximately 130, 10, 3, 2, 1°C/s. Examples neurons are the same as presented in main Fig. 2g.
**b**, Example traces of a pIC broadly tuned neuron responding to the same stimuli used in **a**.

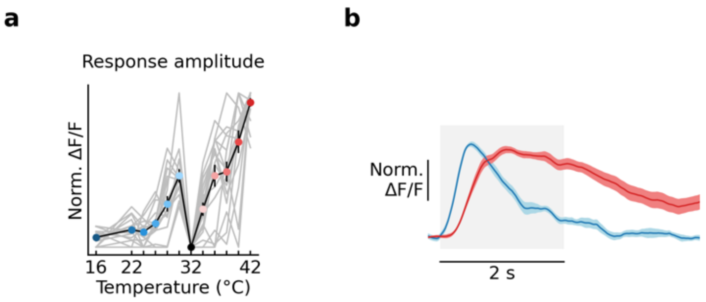


**Supplementary Figure 5. Thermometer cells**

**a**, Response amplitude as a function of stimulus value for neurons with similar response profile to neuron #5 of Fig. 4c. Summary of response amplitude plotted as a function of the thermal stimulus for neurons with similar response profile to neuron #5 of Fig. 4c (n = 16 thermometer neurons out of 411 warm neurons; ~5%). Grey lines show individual neurons, colored filled circles show mean ± s.e.m. at AT 32°C. Most of thermometer neurons are characterized by low threshold response to warming stimuli and had cool responses only for small amplitude cooling stimuli.

**b**, Grand average responses (mean ± s.e.m.) to 2°C cooling (32-30°C) or 10°C warming (32-42°C) from AT 32°C. Thermometer neurons conserve the latency difference observed for cool and warm responses in tuned neurons (Fig. 3b,c). The onset delays of the thermal responses of the thermometers cells indicate that they receive the same input as the tuned and standard broadly tuned neurons. The lack of responses to large cooling stimuli might be explain by disynaptic cortical inhibition (feedback inhibition) driven by a secondary cool channel preferentially and strongly activated by high amplitude cooling stimuli (eg. 32-24°C). These cells highlight complex thermal features possibly constructed by the pIC.

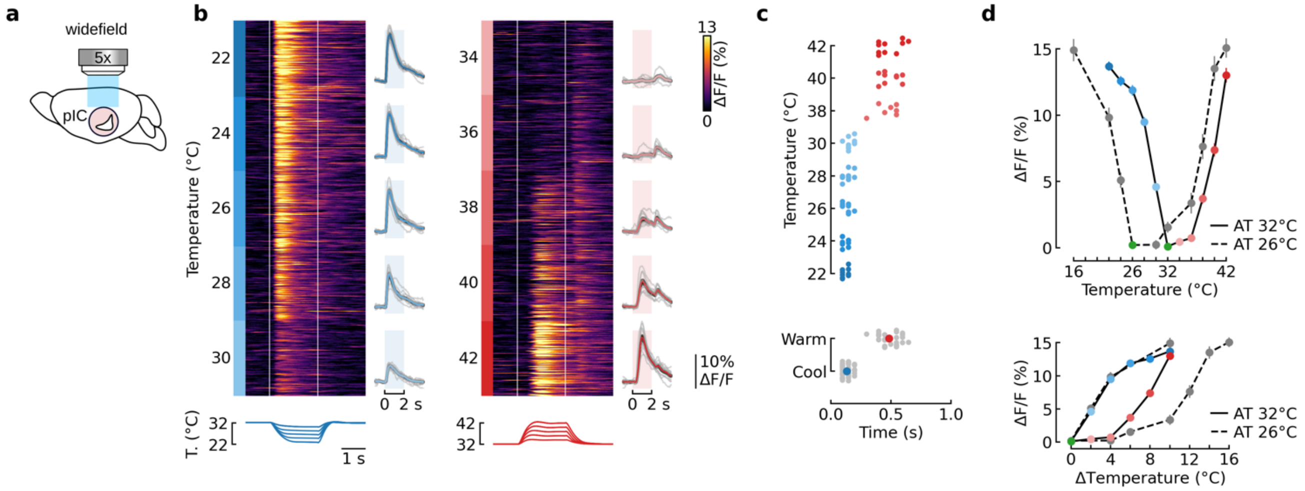


**Supplementary Figure 6. Widefield imaging analysis of thermal amplitude encoding**

**a**, Left, schematic of widefield calcium imaging in a mouse implanted with a cranial window in top of pIC.

**b,** Widefield responses of pIC to forepaw thermal stimulation with different amplitudes (*n* = 123-128 trials per temperature, 10 mice). Heat maps showing widefield responses to thermal stimuli with each line representing a single trial response and thermal stimuli below with white lines showing the onset and the end of the plateau phase of thermal stimuli; colored lines to the right of heatmap show mean ± s.e.m. across trials from all mice and gray lines show mean response from individual mice; colored shaded boxes indicate start of thermal stimulus and end of plateau phase. Right, same as left panels but for warm responses.

**c**, Top, response latencies to different amplitudes of thermal stimuli from individual mice. Bottom, gray filled circles show data from all amplitudes (*n* = 10 mice). Filled circles show mean ± s.e.m.

**d**, Top, peak population response to different amplitude stimuli from experiments presented in **b**. Colored filled circles with solid lines show data with adapted temperature (AT) of 32°C. Gray filled circles with dashed lines show data using AT of 26°C (*n* = 6 mice). Green shows AT. Bottom, same data as in top but showing population response amplitude plotted against thermal stimulus amplitude. Graph shows shift in warm response amplitude but not for cool when presented with a lower AT.

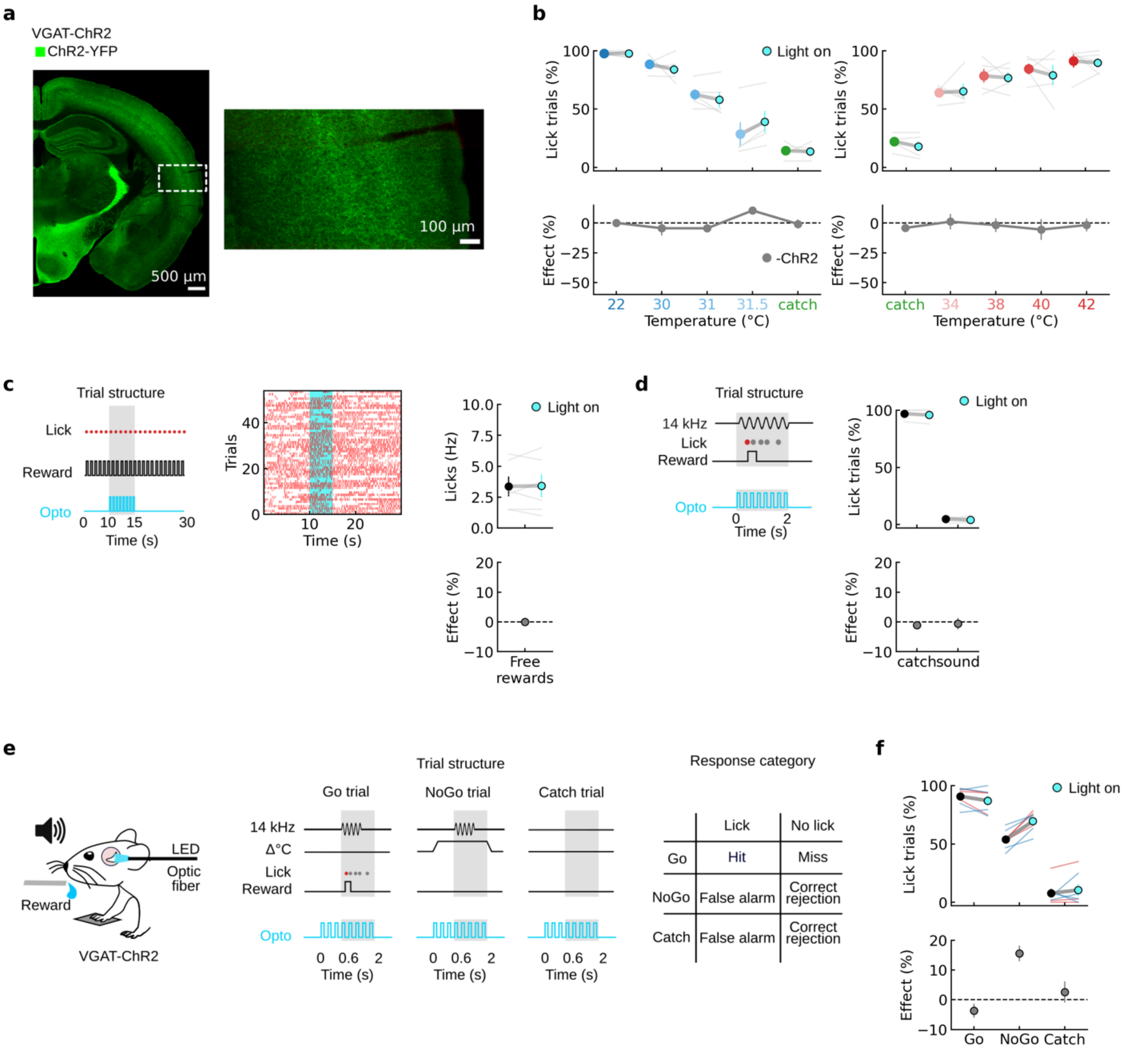


**Supplementary Figure 7. Behavioral control experiments**

**a**, Example coronal slices from VGAT-ChR2 mice showing that expression of ChR2 in pIC is homogeneous and similar to other cortical regions (labelled with anti-GFP antibody).

**b**, Optical stimulation of the thermal zone of pIC in mice that do not express ChR2 (Thy1-GCaMP6s) does not affect cool or warm perception. Behavioral task was identical to that presented in Fig. 5. Top graph shows number of trials with a lick during optical stimulation (cyan) vs. without optical stimulation (blue or red) at different thermal stimulus amplitudes. Gray lines show individual mice ($n$ = 5), colored filled circles are mean ± s.e.m. Lower panel shows the effect of light stimulus (change in percentage of trials with licks, mean ± s.e.m) for the corresponding stimulus amplitude above.

**c**, Optical inhibition of pIC in VGAT-ChR2 mice does not affect spontaneous licking behavior. Left, trial structure showing optical stimulation during free licking, gray bar shows time window for lick rate measurement. Middle, example raster plot of free licking rates in a VGAT-ChR2 mouse during optical stimulation. Right, mean lick rates in VGAT-ChR2 mice during optical stimulation Off (black; 5 s before light stimulus) vs. On (cyan; 5 s during light stimulus) trials. Filled circles show individual mice ($n$ = 6).

**d**, Auditory perception is not altered by optogenetic inhibition of pIC. Task design is same as thermal task but with an acoustic stimulus (14 kHz, 65 dB SPL). Left, structure of optical stimulation trial. Right, mean lick rates in VGAT-ChR2 mice during trials with optical stimulation during the auditory stimuli (cyan) vs. trials without optical stimulation (black). Filled circles show individual mice ($n$ = 4).

**e**, Left, schematic of discrimination task and placement of the optic fiber. Middle, example trial structure showing timing of reward window (gray) and the timing of an optical stimulus during trials with optogenetic manipulation. Filled circles show licks, first lick colored to show rewarded lick. VGAT-ChR2 mice were trained to discriminate a 14 kHz acoustic stimulus (Go trial) from the same stimulus presented 0.6 s after the beginning of a 2 s thermal stimulus (either cooling or warming) (NoGo trial). Right, response categories of the task. Only Go trial was rewarded, NoGo and catch trials (no stimuli) were not rewarded. Training was continued until Go trial hit rate > 70%, NoGo false alarm rate ~50% and false alarm rate < 30%. A criterion of ~50% for the NoGo false alarm rate was chosen to be able to observe an increase or decrease in lick rates during optogenetic inhibition. The amplitude of the thermal stimuli (cooling or warming) was adjusted from mouse to mouse to obtain such criterion.
**f**, Top, proportion of trials with licks in VGAT-ChR2 mice during optical stimulation (cyan) trials vs. without optical stimulation (black). Blue and red lines show experiments with mice trained with either cooling ($n$ = 4) or warming ($n$ = 3) stimuli respectively. Colored filled circles show mean ± s.e.m. Bottom, shows the effect of light stimulus (change in percentage of trials with licks, mean ± s.e.m.). The increase, rather than decrease, in NoGo false alarm rate during optogenetic manipulation clearly indicates the direct and selective involvement of pIC in thermosensory processing.
